# ERBB4 coordinates fibroblast and myeloid cell function during neonatal cardiac regeneration

**DOI:** 10.64898/2026.07.29.741639

**Authors:** Siel Van den Bogaert, Sander Eens, Celine Civati, Eulalie Van den Bogaert, Bo Goovaerts, Julie Cools, Vincent FM Segers, Gilles W De Keulenaer

## Abstract

**BACKGROUND:** Neuregulin-1 (NRG1) is indispensable for scarless regeneration of the injured neonatal mammalian heart, by stimulating cardiomyocyte proliferation through ERBB4 tyrosine kinase receptors. The role of ERBB4 signaling in fibroblasts and myeloid cells in this process remains poorly understood.

**HYPOTHESIS:** We hypothesized that fibroblast– or myeloid-specific *Erbb4* deletion impairs neonatal cardiac regeneration following myocardial infarction (MI).

**METHODS:** Wild type (WT), fibroblast-specific (FB-*Erbb4*-KO) and myeloid-specific (M-*Erbb4*-KO) mice underwent LAD ligation at postnatal day 1 (total n=200). Hearts were harvested at 4, 7, 10, and 21 days post-injury (dpi) for histological analyses, bulk RNA sequencing, and targeted qPCR of the apical LV region. Cardiomyocyte cell-cycle activity was quantified by pH3 and Aurora B kinase staining. Scar size was quantified by Masson’s trichrome staining.

**RESULTS:** FB-*Erbb4*-KO and M-*Erbb4*-KO showed similar levels of cardiomyocyte cell cycle activity post-MI compared to WT controls. However, there were significant differences in the dynamics of myocardial scar size, capillary density and myocardial gene expression. In FB-*Erbb4*-KO mice, infarct scar size at 4 dpi was comparable to WT. Although scar size decreased 6-fold in both WT and KO mice by 7 dpi, FB-*Erbb4*-KO mice retained a significantly larger infarct scar than WT controls. At 10 dpi, KO mice had a larger scar size, lower myocardial capillary density and increased expression of *Col1a1* mRNA levels in the ventricular apex. At 21 dpi, regeneration was nearly complete in both WT and FB-*Erbb4*-KO mice, although the limited residual scar in KO mice was significantly larger compared to WT. In M-*Erbb4*-KO mice, infarct scar size at 4 dpi was 2-fold larger compared to WT. Infarct size remained larger at 7 dpi and 10 dpi while *Mmp2* and *Mmp9* mRNA expression was upregulated. At 21 dpi, regeneration was nearly complete in both WT and KO mice, with no significant size difference between the remaining scars.

**CONCLUSION:** Fibroblast-specific ERBB4 deletion modestly impaired scar resolution, whereas myeloid-specific ERBB4 deletion resulted in a larger initial injury response but did not impair the ultimate regenerative outcome. These findings indicate that ERBB4 in fibroblasts and myeloid cells coordinates regenerative microenvironment dynamics beyond cardiomyocyte proliferation rather than being an indispensable switch for neonatal cardiac regeneration.

## Introduction

Cardiac regeneration after injury is limited in adult mammals, primarily because cardiomyocytes exit the cell cycle shortly after birth (1–3). This loss of proliferative capacity results in the formation of fibrotic scar tissue and minimal repair of functional myocardium after injury. In contrast, the neonatal heart retains a transient regenerative potential during a developmental window in which cardiomyocytes retain proliferative competence (1,4). Pioneering studies demonstrated that mice injured on postnatal day 1 (P1) can fully regenerate cardiac tissue after apical resection or myocardial infarction (MI) (1,2). However, this regenerative capacity declines rapidly and is largely lost by postnatal day 7 (P7), beyond which fibrotic scar formation predominates over myocardial tissue replacement, resulting in limited functional recovery (1,5). This postnatal loss of regenerative capacity has been associated with cardiomyocyte cell-cycle arrest, metabolic adaptation, increased oxidative stress, and functional maturation of the postnatal heart. Collectively, these observations suggest that the transition from regeneration to fibrotic repair reflects not only intrinsic changes in cardiomyocytes but also broader alterations in the cardiac cellular microenvironment (5,6).

Successful regenerative repair of the neonatal heart requires tight coordination of cardiomyocyte proliferation, rapid revascularization, extracellular matrix remodeling, and a balanced inflammatory response (7,8). These processes are orchestrated by multiple signaling pathways, among which neuregulin-1 (NRG1)-ERBB signaling has emerged as an important contributor to cardiac regeneration (9–11). Upon NRG1 binding, ERBB4 undergoes homo– or heterodimerization, most commonly with ERBB2, resulting in activation of downstream signaling pathways including PI3K-AKT and MAPK/ERK (12). Thus far, the regenerative effects of NRG1-ERBB4 signaling have been studied predominantly in cardiomyocytes. In these cells, NRG1 stimulates proliferation through ERBB4 activation, whereas genetic inactivation of *Erbb4* reduces cardiomyocyte cell-cycle activity and impairs cardiac development (10,12,13). Furthermore, the regenerative response to NRG1 is restricted to a narrow neonatal window, during which progressive postnatal downregulation of ERBB2 limits cardiomyocyte proliferative responsiveness after the first week of life (9,14,15). A similar developmental window has been described in human infant myocardium, where the loss of responsiveness to NRG1 coincides with cardiomyocyte maturation (13,14).

Despite the well-established role of ERBB4 signaling in cardiomyocytes, its contribution in non-cardiomyocyte cell types involved in neonatal cardiac regeneration remains poorly understood. This is particularly important because neonatal cardiac regeneration depends on coordinated interactions between cardiomyocytes and other cardiac cell types, including fibroblasts, endothelial cells, and immune cells (7,16,17). Among these, fibroblasts and immune cells play central roles in extracellular matrix remodeling and inflammatory responses that are essential for scarless repair of the heart. Emerging evidence indicates that fibroblasts also engage ERBB4 signaling during cardiac injury. In adult mice, ERBB4 activation has been associated with reduced fibroblast proliferation and attenuated myocardial fibrosis, suggesting a potential role in regulating post-injury tissue remodeling (18–20). Immune cells, including macrophages, also express ERBB4 which has been implicated in modulating inflammatory responses and cellular activation following injury (19,21). Following MI, immune cells are rapidly recruited to the damaged myocardium, initially adopting a pro-inflammatory phenotype with corresponding cytokine release and clearance of necrotic tissue. As repair progresses, these cells transition toward a reparative state promoting wound healing and constraining fibrosis (7,22–24). Together, these observations suggest that ERBB4 signaling may influence neonatal cardiac regeneration through regulation of the regenerative microenvironment in addition to its established effects on cardiomyocyte proliferation.

This study aims to expand the current framework of ERBB4 regenerative biology beyond cardiomyocytes by defining its role in fibroblasts and myeloid cells during neonatal cardiac regeneration. Investigating ERBB4 function in these non-cardiomyocyte cell populations may provide new insights into the cellular mechanisms driving neonatal cardiac repair. We hypothesized that selective disruption of NRG1-ERBB4 signaling in fibroblasts and myeloid cells, while preserving cardiomyocyte ERBB4 activity, will impair the efficiency of neonatal cardiac regeneration, resulting in delayed scar resolution and persistent residual fibrosis.

## Methods

### Animals

The Ethical Committee for Animal Testing of the University of Antwerp granted approval for all procedures (approval no. 2022-56), conform to the EU Directive (2010/63) on the protection of animals used for scientific purposes. Experiments were reported in accordance with the ARRIVE guidelines. Mice were housed in the central animal care facility of Antwerp University under standard conditions (20-24°C, 45-65% relative humidity, 12:12 h light/dark cycle) with free access to regular chow and water.

Fibroblast-specific *Erbb4* gene deletion: mice with fibroblast-specific deletion of *Erbb4* were generated by crossing fibroblast-specific *Col1a2*-Cre recombinase mice (*Tg(Col1a2-cre)23Angl*, kindly provided by Dr. P. Angel, German Cancer Research Center (Das Deutsche Krebsforschungszentrum, DKFZ)) with *Erbb4^f/f^* mice (*B6;129-Erbb4tm1Fej/Mmucd*,#010439-UCD, MMRRC). Because homozygous fibroblast-specific deletion was not viable, hemizygous *Erbb4^f/+^ Col1a2^+^* mice were used for all experiments. *Erbb4^+/+^ Col1a2^+^* and wild-type (WT) littermate controls were used.

Myeloid-specific *Erbb4* gene deletion: myeloid-specific *Erbb4^f/f^ LysM-Cre^+/-^* knock-out (KO) mice were generated by crossing myeloid-specific *LysM-Cre^+/+^* mice (B6.129P2-*Lyz2tm1(cre)lfo/J*, 004781, JAX) with *Erbb4^f/f^* mice. *Erbb4^+/+^ LysM-Cre^+/-^* WT littermates served as controls.

For both mouse models, genotyping was performed after completion of the experimental procedures to ensure investigator blinding.

To confirm *Erbb4* deletion, *Erbb4* mRNA expression was quantified by RT-qPCR in isolated cardiac fibroblasts and bone marrow-derived macrophages from WT and KO mice. RNA was isolated from both isolated cell types with (NucleoSpin RNA XS Macherey Nagel, 740902) according to the manufacturer’s instructions. Relative RNA expression was quantified by performing two-step RT-qPCR using the TaqMan^TM^ Reverse Transcription Reagents (N8080234, ThermoFisher, Waltham, MA, USA), TaqMan^TM^ Universal PCR Master Mix (4304437, ThermoFisher, USA) and Taqman^TM^ primers. RT-qPCR was performed using the QuantStudio^TM^ 3 instrument (ThermoFisher, USA). All data were normalized against housekeeping genes *Gapdh* and *Rpl32.* Relative gene expression was evaluated according to the 2 (−ΔΔCT) method. Following primers were used: Erbb4 (Mm01256793_m1) Rpl32 (Mm02528467_g1) and Gapdh (Mm99999915_g1).

Protein lysates from the same cell types were analyzed by western blot using antibodies against ERBB4 (Cell signaling, 4795, 1:300) with GAPDH (Cell signaling, 97166, 1:1000) or Vinculin (Cell Signaling, 13901S, 1:5000) as housekeeping for normalization. Validation of *Erbb4* deletion at the mRNA and protein level is shown in Figure 1B.

**Figure 1.**
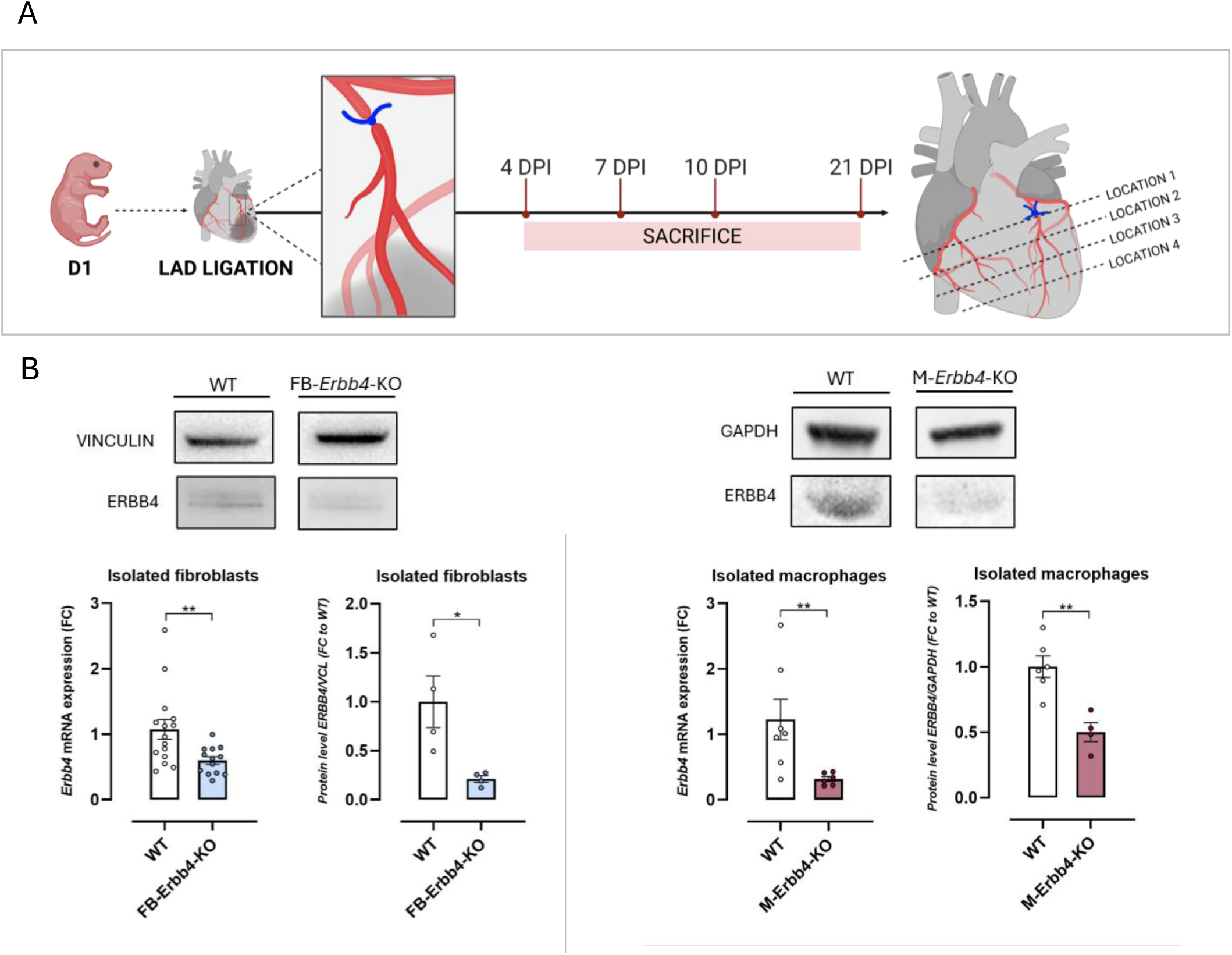
Overview of the study design and genotype validation. **(A)** Experimental design. Myocardial infarction (MI) was induced at postnatal day 1 (P1) by ligation of the left anterior descending (LAD) coronary artery. Hearts were collected at 4, 7, 10, and 21 days post-injury (dpi). Transverse sections spanning the infarct region were obtained at four different locations from the ligation site to the apex for histological analysis. **(B)** Validation of cell type-specific *Erbb4* deletion at the mRNA and protein levels. Cardiac fibroblasts isolated from WT and FB-*Erbb4*-KO hearts and bone marrow-derived macrophages from WT and M-*Erbb4*-KO mice were analyzed by qPCR and Western blotting. Differences between groups were analyzed using unpaired Student’s t-test, mean ± SEM. * p ≤ 0.05, ** p ≤ 0.01.

### Neonatal MI model

Neonatal mice of both sexes (total n=200; female n=112 and male n=88), including transgenic lines and WT littermate controls, were used in this study. At P1, mice were anesthetized by hypothermia on an ice bed for 3-4 min until an adequate level of anesthesia was achieved. Anesthesia was confirmed by cessation of movement and absence of response to toe pinch. Left lateral thoracotomy was performed at the fourth intercostal space by blunt dissection of the intercostal muscles following skin incision. The left anterior descending (LAD) coronary artery was ligated at the mid-ventricular level using a 10-0 Prolene® suture (2794G). Successful ligation was confirmed by immediate blanching of the ventricular wall. After ligation, the thoracic wall and skin were closed using an 8-0 non-absorbable Prolene® suture (F1894). Sham-operated mice underwent the same procedure, including hypothermic anesthesia and thoracotomy, but without LAD ligation. After surgery, pups were gently cleaned and allowed to recover in a heated chamber (37 °C) with abundant bedding from the dam’s cage. Once all littermates were adequately recovered, they were returned to their dam for the duration of the follow-up period. Pups were monitored to confirm postoperative recovery and maternal acceptance (1,25,26).

### Sample collection

At the end of the follow-up period (4, 7, 10, or 21 dpi), mice were weighed, anesthetized with isoflurane, and euthanized by decapitation (4-10 dpi) or cervical dislocation (21 dpi). The hearts were collected and weighed. For histological and transcriptomic analyses, cardiac tissue distal to the LAD ligation site was collected. Owing to the limited size of the neonatal heart, separate cohorts were used for each analysis. For histology with subsequent formalin-fixed paraffin-embedded (FFPE) RNA isolation, transverse cardiac cross-sections and the apical tissue were used (n=6-16 per genotype per timepoint), whereas for bulk RNA sequencing the entire cardiac tissue distal to the ligation site was used (n=5 per group). Heart samples for histology were fixed and paraffin embedded. Samples for transcriptomics were snap-frozen in liquid nitrogen and stored at –80 °C until analysis. Following sample collection, the sex of the pups was confirmed by gonadal examination.

Figure 1A provides an overview of the study design, which was applied to both genotypes (FB-*Erbb4*-KO, M-*Erbb4*-KO) and their respective littermate controls.

### Histological analysis and immunohistochemistry

Heart samples were fixed in 4% paraformaldehyde (PFA), paraffin-embedded, and sectioned at 5 µm thickness. Transverse sections were obtained at four distinct locations between the ligation site and the apex (Figure 1A). Sections were stained with Masson’s Trichrome according to standard protocols for infarct scar measurement (25,26). Total scar size was quantified as the ratio of scar area to total left ventricular area for each section. Values from all sections were combined to determine total infarct size per heart. For immunohistochemistry, sections were incubated overnight at room temperature with specific primary antibodies for cardiac Troponin T (Anti-cardiac troponin T (1C11) (cTnT), Abcam ab8295, 1:200), phospho-histone 3 (Anti-Histone H3 (phospho S10) (pH3), Abcam ab47297, 1:100) and Aurora B kinase (Anti-Aurora B, Abcam ab2254, 1:200). The following day, the sections were washed three times with PBS and incubated for 1 h with the corresponding secondary antibody, conjugated to AlexaFluor™ 488 (cTNT) or AlexaFluor™ 546 (pH3 and Aurora B) (1:400, Invitrogen). Nuclei were counterstained with DAPI-containing mounting medium (Vector Laboratories, H-1500). Cardiomyocyte hypertrophy and vascularization were assessed by isolectin-B4 (AF 488, ThermoFisher scientific, I21411, 1:50) / WGA (AF594, ThermoFisher scientific, W11262, 1:100) / DAPI staining. Images were acquired using Olympus BX43 microscope or CELENA^®^ S Digital Imaging System and analyzed with ImageJ v2.14.0 software. Image quantifications were performed by an investigator blinded to genotype and intervention.

### Transcriptomics

Snap-frozen cardiac samples were processed for RNA isolation and RNA sequencing (outsourced to Azenta Life Sciences, Germany). Total RNA was isolated, and RNA quality and quantity were assessed using standard quality control procedures. Sequencing libraries were prepared using a stranded mRNA library preparation protocol with poly(A) selection. Libraries were sequenced on an Illumina NovaSeq 6000 platform (paired-end, 2×150 bp), yielding approximately 20 million reads per sample. Raw sequencing data were processed by Azenta, including read alignment to the mouse reference genome (mm10) using STAR and generation of gene-level count matrices.

The gene count matrix was imported into Omics Playground (BigOmics Analytics (27)) for differential gene expression analysis, using limma, edgeR and DESeq2 statistical frameworks. To minimize method-specific bias, differentially expressed genes were identified using a conservative meta.q statistic, defined as the highest false discovery rate (FDR)-adjusted P value obtained across the three methods.

Gene set enrichment analysis (GSEA) was subsequently performed in R (v4.4.1). Gene Ontology (GO) Biological Process enrichment analysis was performed using the clusterProfiler package with gene annotation provided by org.Mm.eg.db. Hallmark gene set enrichment analysis was performed using the fgsea package with Hallmark gene sets obtained from the Molecular Signatures Database (MSigDB). Gene sets with an FDR-adjusted P value <0.05 were considered significantly enriched.

### FFPE RNA isolation and RT-qPCR analysis

Total RNA was isolated from FFPE heart tissue sections of the most apical location of the heart (location 4, see Figure 1A) using a commercially available kit (Macherey Nagel 740969) according to the manufacturer’s instructions. RNA concentration and quality were assessed using NanoDrop ND-1000 spectrophotometer. Purified RNA was used for downstream gene expression analyses. Relative RNA expression was quantified by performing two-step RT-qPCR using the TaqMan^TM^ Reverse Transcription Reagents (N8080234, ThermoFisher, Waltham, MA, USA), TaqMan^TM^ Universal PCR Master Mix (4304437, ThermoFisher, USA) and Taqman^TM^ primers (see below). RT-qPCR was performed using the QuantStudio^TM^ 3 instrument (ThermoFisher, USA). All data were normalized against housekeeping genes *Rpl32* and *Gapdh*. Relative gene expression was evaluated according to the 2 (−ΔΔCT) method.

Following primers were used: Rpl32 (Mm02528467_g1), Gapdh (Mm99999915_g1), *Col1a1* (Mm008001666_g1), *Mmp2* (Mm00439498_m1), *Mmp9* (Mm00442991_m1) and *Timp1* (Mm00441818_m1).

### Statistical analysis

Normality was assessed through visual inspection (i.e., QQ plot) and a Shapiro-Wilk test or Kolmogorov-Smirnov test. For immunohistochemistry and RT-qPCR analyses, differences between groups were analyzed with a two-way ANOVA followed by a Tukey’s multiple-comparisons test (normal distribution) or Kruskal-Wallis test followed by Dunn’s multiple-comparisons test (non-normal distribution). KO validation and scar size comparisons were analyzed using an unpaired Student’s t-test for normally distributed data or the Mann-Whitney U test for non-normally distributed data. Differences were considered statistically significant when p ≤ 0.05. Data are expressed as the mean ± standard error of the mean (SEM). Statistical analyses were performed and graphs were generated using GraphPad Prism v9.3.1.

## Results

### Effect of fibroblast– and myeloid-specific *Erbb4* deletion on post-MI cardiomyocyte cell-cycle activity, cardiomyocyte cross-sectional area, and myocardial capillary density

Cardiac regeneration after MI in neonatal mice is characterized by cardiomyocyte proliferation, limited hypertrophic remodeling, and rapid revascularization of the injured myocardium (6,13). To determine whether fibroblast-specific or myeloid-specific *Erbb4* gene deletion affects these hallmark features of neonatal cardiac regeneration, cardiomyocyte mitotic activity, cardiomyocyte cross-sectional area, and capillary density were quantified.

Cardiomyocyte cell-cycle activity was assessed by pH3 and Aurora B kinase staining in cTnT positive cells. Analyses were performed at 4 and 7 dpi, corresponding to the peak regenerative window in the neonatal mouse heart (1). MI significantly increased cardiomyocyte cell-cycle activity in cTnT positive cells compared to sham-operated controls at both time points. However, no significant differences were observed between WT mice and KO mice in either the FB-*Erbb4*-KO or M-*Erbb4*-KO model at 4 or 7 dpi (Figure 2A).

**Figure 2.**
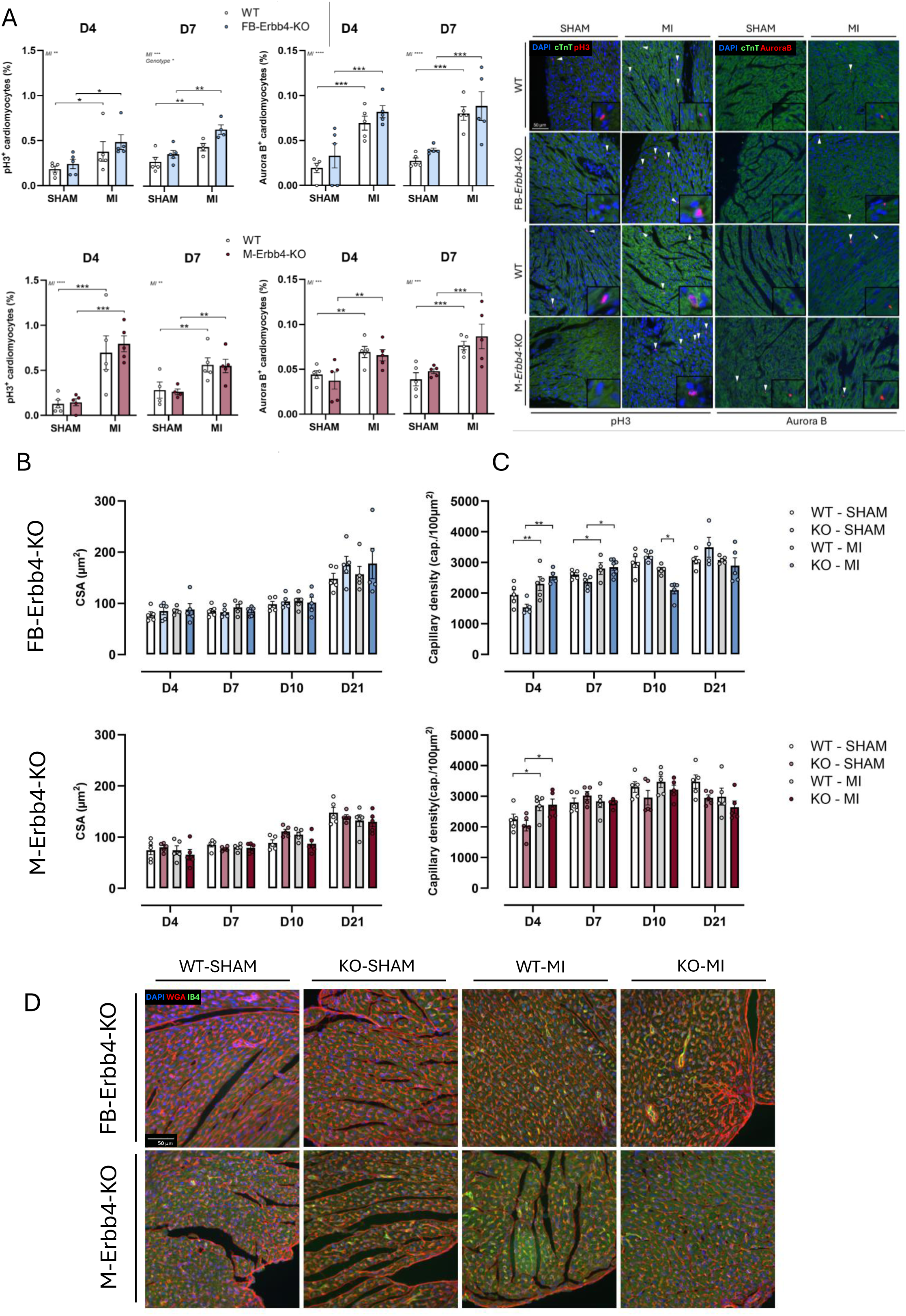
Cell type-specific deletion of *Erbb4* does not affect cardiomyocyte proliferation or hypertrophy but transiently alters capillary remodeling after MI. **(A)** Quantification of cardiomyocyte cell-cycle activity by phospho-histone H3 (pH3) and Aurora B kinase staining in cardiac troponin T (cTnT)-positive cardiomyocytes at 4 and 7 dpi. FB-*Erbb4*-KO and M-*Erbb4*-KO mice were compared to wild-type (WT) littermates under sham and myocardial infarction (MI) conditions. Representative immunofluorescence images are shown (cTnT, green; pH3/AuroraB, red; nuclei, DAPI in blue). **(B)** Quantification of cardiomyocyte cross-sectional area (CSA) at 4, 7, 10, and 21 dpi in FB-*Erbb4*-KO and M-*Erbb4*-KO mice and WT littermates following sham surgery or MI. **(C)** Quantification of myocardial capillary density (capillaries/mm²) by isolectin B4 staining at 4, 7, 10, and 21 dpi. **(D)** Representative immunofluorescence images of myocardial sections at 10 dpi showing capillary networks (isolectin B4, green) and cell boundaries (WGA, red) in WT and *Erbb4* KO mice under sham and MI conditions. Data are presented as mean±SEM. Each dot represents an individual animal. Statistical significance was determined using 2-way ANOVA with multiple comparisons correction. * p ≤ 0.05, ** p ≤ 0.01, *** p ≤ 0.001

Cardiomyocyte cross-sectional area was quantified to evaluate compensatory hypertrophic growth up to 21 dpi. No significant differences in cardiomyocyte cross-sectional area were observed between KO and WT mice in either transgenic line, under sham or after LAD ligation (Figure 2B, D).

Capillary density was next quantified by isolectin B4 staining (Figure 2C, D). At 4 and 7 dpi, MI induced a significant increase in capillary density compared to sham-operated controls, independently of genotype. At 10 dpi, FB-*Erbb4*-KO mice, but not M-*Erbb4*-KO mice, showed a transient reduction in capillary density compared to WT controls (p=0.0157), potentially suggesting a deficient maintenance or maturation of capillaries during the remodeling phase in FB-*Erbb4*-KO hearts.

### Effect of fibroblast– and myeloid-specific *Errb4* deletion on post-MI scar size in the regenerating neonatal heart

In FB-*Erbb4*-KO mice, infarct scar size at 4 dpi was comparable to that of WT controls, indicating that initial scar formation was not affected by the genotype. By 7 dpi, scar size had markedly declined in both groups, showing an approximately sixfold reduction compared with 4 dpi, indicative of a robust regenerative response during the early postnatal period. Nevertheless, a significant difference was observed between genotypes, with FB-*Erbb4*-KO mice displaying a larger residual scar compared to WT littermates (0.3% vs 0.6%, p=0.0275). This difference persisted at 10 dpi, with a larger residual scar in FB-*Erbb4*-KO mice compared to WT mice (0.1% vs 0.6%, p=0.0499). At 21 dpi, scar resolution was nearly complete in both genotypes, although FB-*Erbb4*-KO mice retained a significantly larger residual scar than WT controls (0.08% vs 0.2%; p=0.0033) (Figure 3B, C).

**Figure 3.**
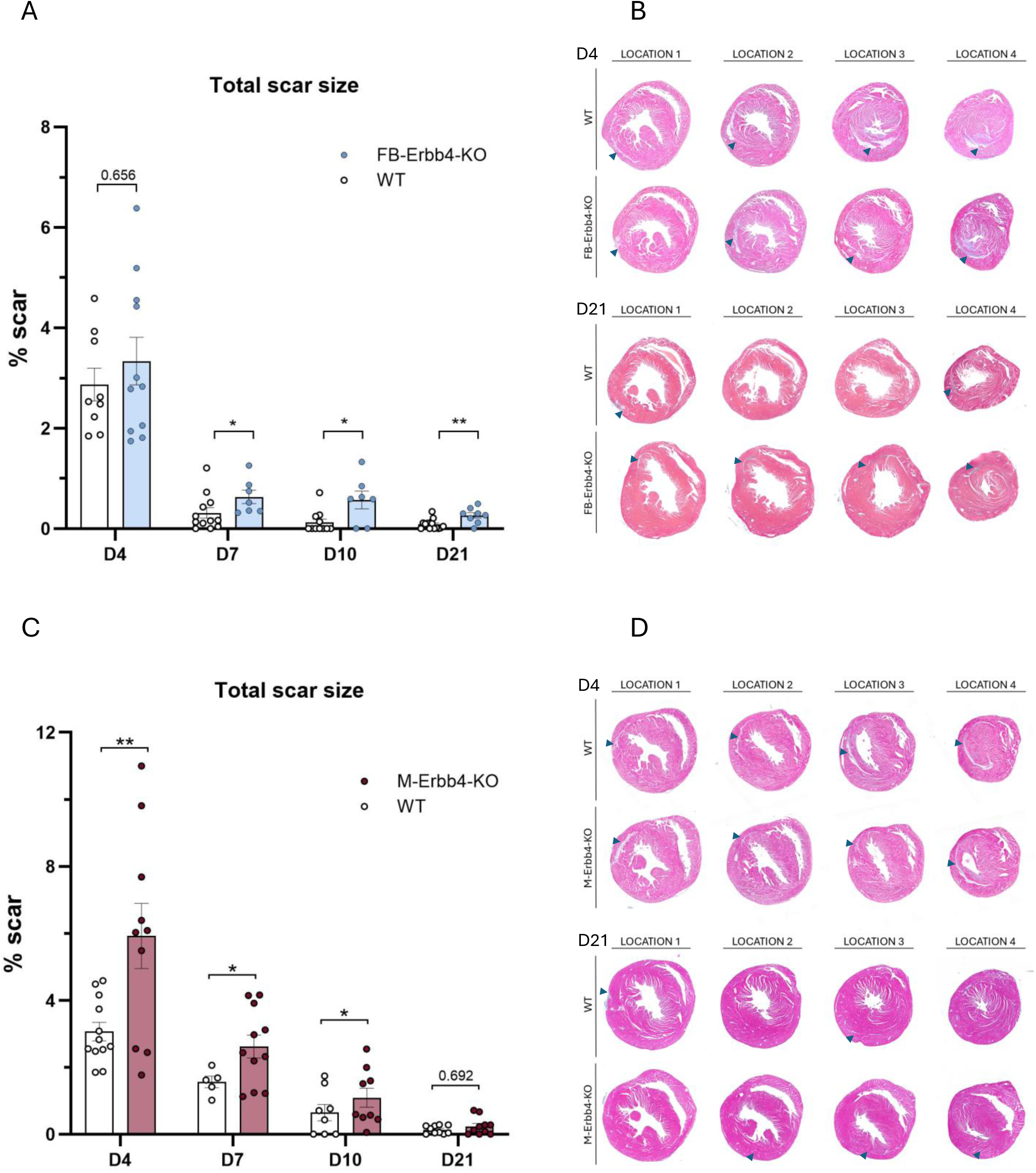
Cell type-specific deletion of *Erbb4* differentially regulates infarct resolution during neonatal cardiac regeneration. **(A)** Quantification of total scar size (% of ventricular area) in FB-*Erbb4*-KO mice and WT littermates at the indicated time points after MI. **(B)** Representative Masson’s trichrome-stained transverse heart sections from WT and FB-*Erbb4*-KO mice at 4 and 21 dpi, shown across locations 1-4. Arrows indicate the infarcted region. **(D)** Quantification of total scar size in M-*Erbb4*-KO and WT littermates at the indicated time points after MI. **(E)** Representative Masson’s trichrome-stained transverse heart sections from WT and M-*Erbb4*-KO mice at 4 and 21 dpi, shown across locations 1-4. Arrows indicate the infarcted region. Data are presented as mean±SEM. Each dot represents an individual animal. Statistical significance was determined using unpaired Student’s t-test or Mann Whitney test. *p ≤ 0.05, **p ≤ 0.01, *** p ≤ 0.001

In M-*Erbb4*-KO mice, infarct scar size at 4 dpi was significantly larger compared to WT controls (3.1% vs 5.9%, p=0.0062), suggesting an enhanced early response to infarction. Scar size subsequently declined in both genotypes but remained significantly larger in myeloid-specific *Erbb4* KO mice at 7 dpi (1.6% vs 2.6%, p=0.0499) and 10 dpi (0.6% vs 1.1%, p=0.0232). By 21 dpi, infarct resolution was nearly complete in both groups, with no significant difference in residual scar size between genotypes (Figure 3D, E).

### Effect of fibroblast-specific *Erbb4* deletion on post-MI cardiac gene expression

To determine the effect of *Erbb4* gene deletion in fibroblasts on the transcriptome of the regenerating neonatal hearts, bulk RNA sequencing was performed on myocardial tissue collected at 10 dpi, corresponding to a critical phase of scar remodeling and resolution in both KO models. Differential expression analysis revealed only a limited number of significantly altered genes between FB-*Erbb4*-KO mice and WT mice, as illustrated in the volcano plot (Figure 4A). In line with these small transcriptional differences, Hallmark analysis did not demonstrate enrichment of canonical inflammatory or extracellular matrix remodeling pathways (Figure 4B).

**Figure 4.**
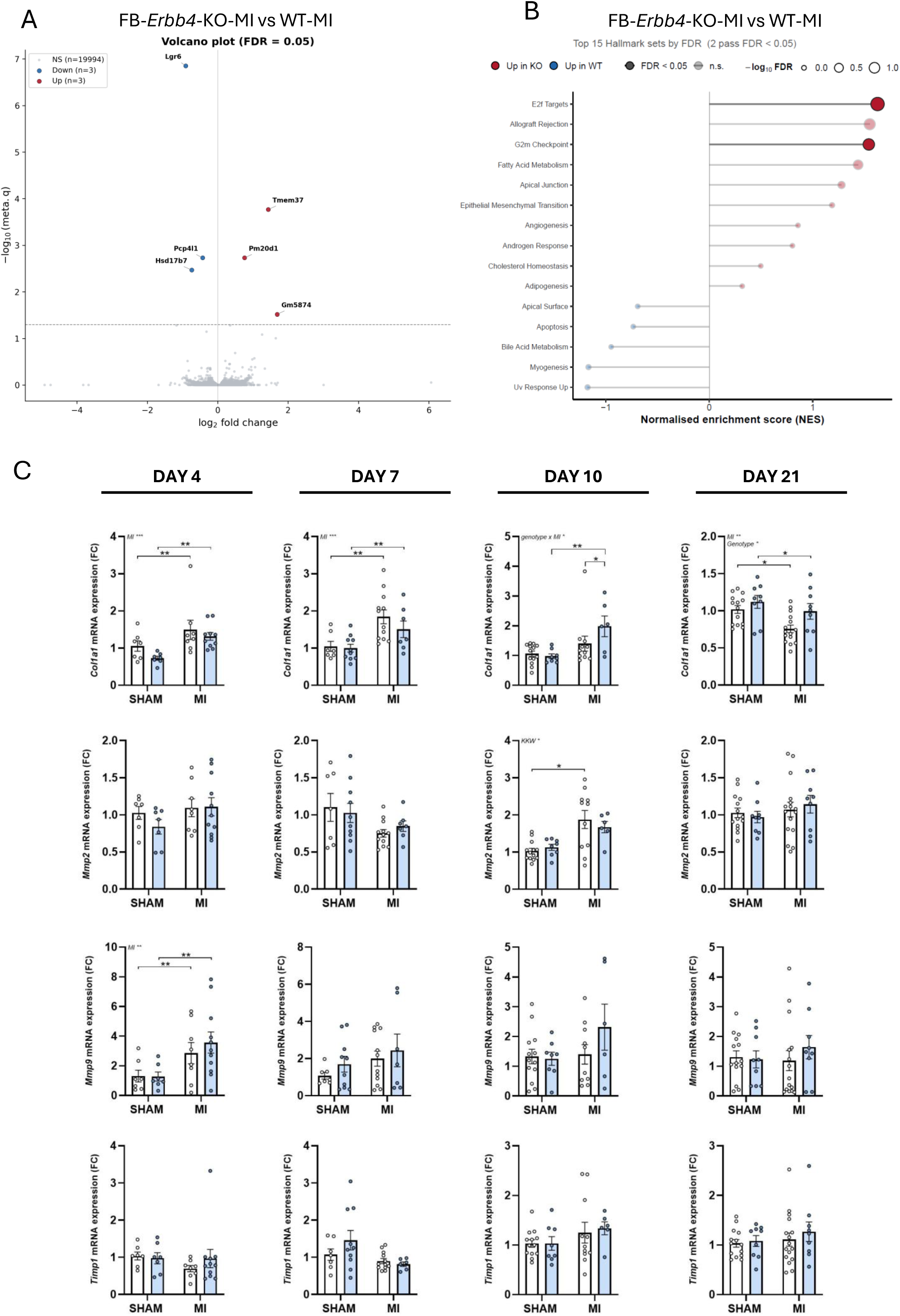
Fibroblast-specific *Erbb4* deletion induces modest transcriptional changes associated with extracellular matrix remodeling during neonatal cardiac regeneration. **(A)** Volcano plot showing differential gene expression in myocardial tissue from FB-*Erbb4*-KO and WT mice at 10 dpi, as assessed by bulk RNA sequencing. Differentially expressed genes are highlighted (red, upregulated; blue, downregulated in FB-*Erbb4*-KO vs WT). **(B)** Hallmark gene set enrichment analysis (GSEA) of differentially expressed genes in FB-*Erbb4*-KO versus WT hearts at 10 dpi. Gene sets with an FDR-adjusted P < 0.05 are shown. **(C)** Targeted RT-qPCR analysis of extracellular matrix remodeling-associated genes (*Col1a1*, *Mmp2*, *Mmp9*, *Timp1*) in the infarct region at 4, 7, 10, and 21 dpi in FB-*Erbb4*-KO and WT mice under sham and MI conditions. Gene expression was normalized to housekeeping genes and expressed relative to WT sham controls. Data are presented as mean±SEM. Each dot represents an individual animal. Statistical significance was determined using 2-way ANOVA with multiple comparisons correction. *p ≤ 0.05, **p ≤ 0.01, *** p ≤ 0.001

Because the tissue analyzed for bulk RNA sequencing included the entire left ventricle, changes in gene expression in the infarcted tissue could have been diluted. To address this, a targeted qPCR approach on the mRNA expression of selected fibrotic/remodeling-associated genes (*Col1a1, Mmp2, Mmp9, Timp1*) was performed on the apical section, which mainly consists of infarcted/regenerating tissue after LAD ligation. Figure 4C shows the expression of these genes in WT and FB-*Erbb4*-KO mice following sham and MI at 4,7, 10 and 21 dpi. At 4 dpi and 7 dpi, infarcted hearts showed upregulated expression of *Col1a1* regardless of the genotype. Strikingly at 10 dpi, FB-*Erbb4*-KO mice displayed a significantly increased expression of *Col1a1* compared to WT controls (p=0.0404). No significant changes in gene expression were observed at 21 dpi.

### Effect of myeloid-specific *Erbb4* deletion on post-MI cardiac gene expression

In contrast to the limited transcriptional alterations observed in FB-*Erbb4*-KO hearts, bulk RNA sequencing of M-*Erbb4*-KO hearts at 10 dpi identified 28 upregulated and 37 downregulated genes compared to WT mice (Figure 5A). Among the differentially expressed genes were the fibrotic markers *Emilin2* and *Acta2*, as well as inflammation-associated genes including *Gmfg*, and *Tlr2* (Figure 5A). Hallmark analysis demonstrated broad enrichment of inflammatory and tissue remodeling-associated pathways in M-*Erbb4*-KO hearts, including interferon responses, IL6-JAK-STAT3 signaling, inflammatory response, and epithelial-mesenchymal transition gene sets (Figure 5B). Consistent with these findings, GO analysis identified enrichment of biological processes related to immune cell recruitment, innate and adaptive immune responses, extracellular matrix organization, and regulation of angiogenesis. Together, these transcriptomic changes indicate persistent inflammatory and remodeling activity in the absence of myeloid ERBB4 signaling.

**Figure 5.**
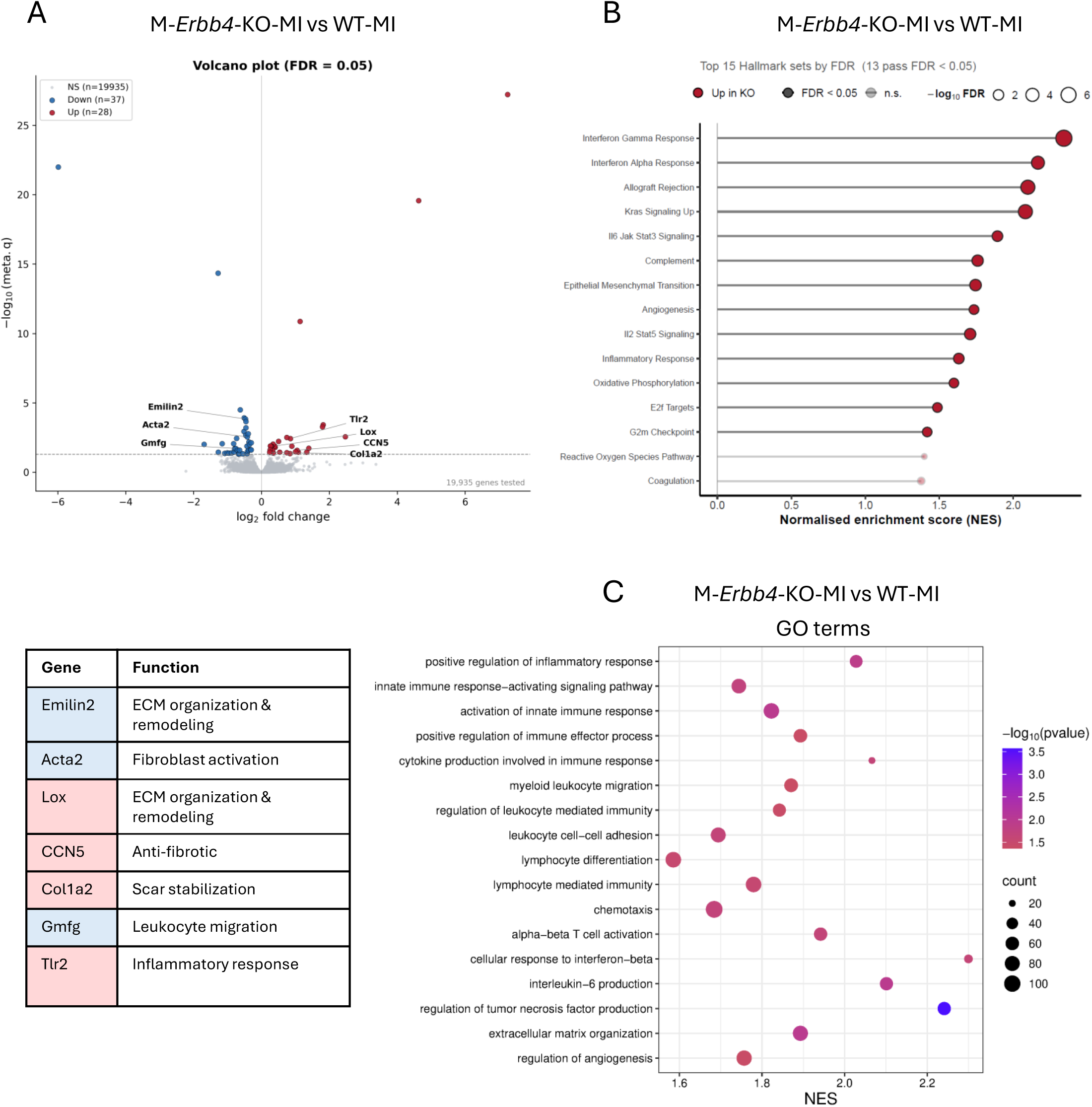
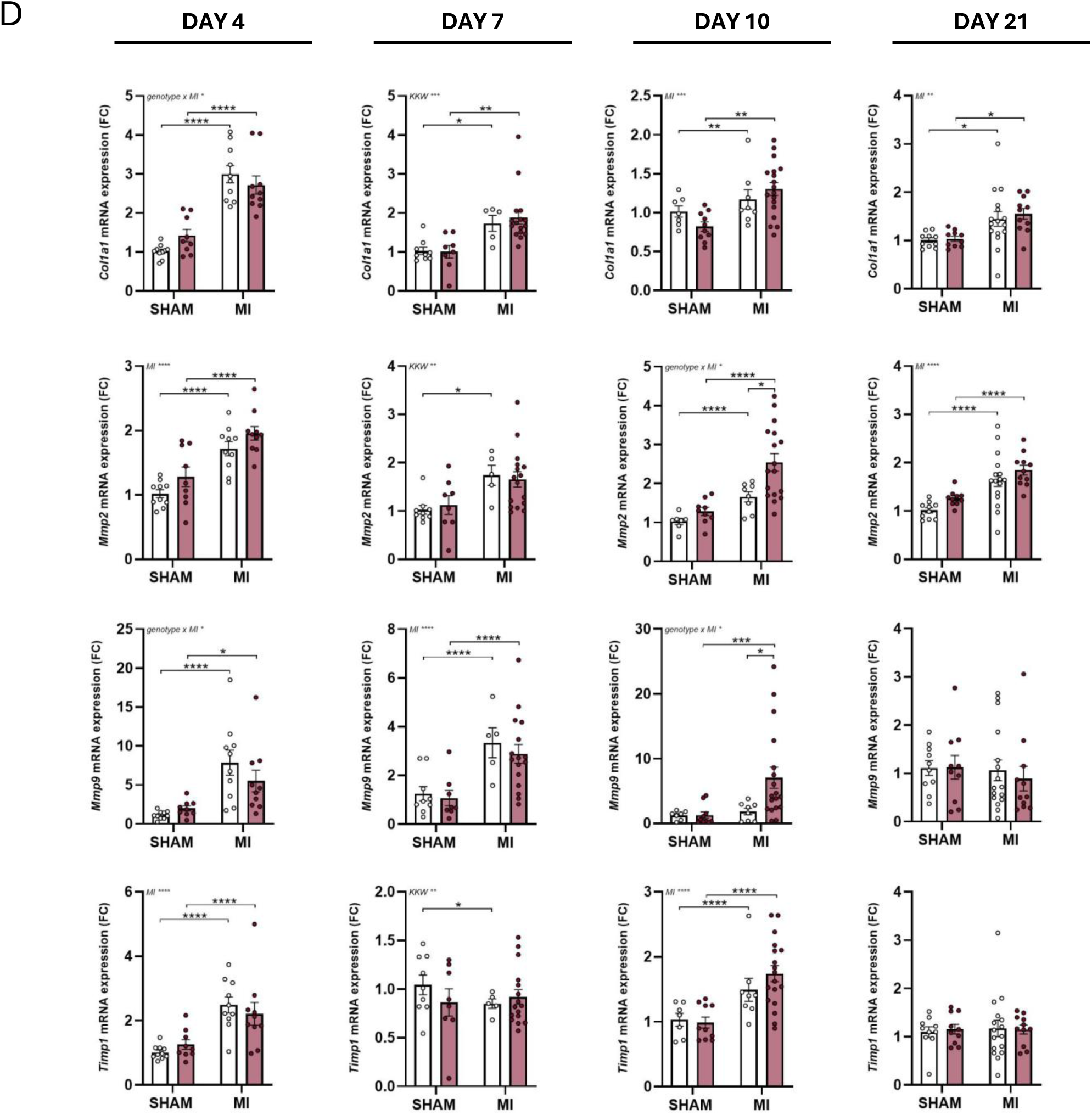
Myeloid-specific *Erbb4* deletion promotes inflammatory and extracellular matrix remodeling programs during neonatal cardiac regeneration. **(A)** Volcano plot showing differential gene expression in myocardial tissue from M-Erbb4-KO and WT mice at 10 dpi, as determined by bulk RNA sequencing. Differentially expressed genes are highlighted (red, upregulated; blue, downregulated in M-Erbb4-KO versus WT), including genes associated with inflammation and extracellular matrix remodeling. The accompanying table summarizes the highlighted genes and their putative biological functions. **(B)** Hallmark gene set enrichment analysis (GSEA) of differentially expressed genes in M-*Erbb4*-KO versus WT hearts at 10 dpi. Gene sets with an FDR-adjusted P < 0.05 are shown. **(C)** Gene Ontology (GO) Biological Process GSEA of differentially expressed genes in M-Erbb4-KO versus WT hearts at 10 dpi. The dot plot shows significantly enriched biological processes related to inflammatory signaling, immune cell recruitment, extracellular matrix organization, and regulation of angiogenesis. Dot size represents the number of genes contributing to each gene set, and color indicates the FDR-adjusted P value. **(D)** Targeted RT-qPCR analysis of extracellular matrix remodeling-associated genes (*Col1a1*, *Mmp2*, *Mmp9*, *Timp1*) in the infarct region at 4, 7, 10, and 21 dpi in M-*Erbb4*-KO and WT mice under sham and MI conditions. Gene expression was normalized to housekeeping genes and expressed relative to WT sham controls. Data are presented as mean±SEM. Each dot represents an individual animal. Statistical significance was determined using 2-way ANOVA with multiple comparisons correction. *p ≤ 0.05, **p ≤ 0.01, *** p ≤ 0.001, **** p ≤ 0.0001

To further characterize these responses over time and more specifically in the infarcted/regenerating zone, a targeted qPCR analysis was performed using the same panel of fibrotic/remodeling-associated genes (*Col1a1, Mmp2, Mmp9, Timp1*) (Figure 5C). At 4 dpi, infarcted hearts showed upregulated expression of all remodeling-associated genes regardless of the genotype. At 7 dpi this pattern remained consistent for WT infarcted hearts with the exception of *Timp1* which was significantly downregulated. In infarcted KO hearts, only *Col1a1* and *Mmp9* remained significantly upregulated. At 10 dpi, KO-MI hearts exhibited significant upregulation of the matrix metalloproteinases *Mmp2* (p=0.0103) and *Mmp9* (p=0.0412) relative to WT-MI controls, consistent with enhanced matrix turnover and remodeling activity. No significant changes in gene expression were observed at 21 dpi.

## Discussion

The neonatal heart possesses a transient regenerative capacity that is restricted to a narrow window (1,5,12). Following myocardial injury, successful neonatal regeneration is characterized by a coordinated sequence of events, including robust cardiomyocyte mitotic activity without compensatory hypertrophic growth, transient scar formation followed by progressive scar resolution, restoration of the microvasculature, and tightly regulated inflammatory and fibrotic responses that resolve as regeneration proceeds. Disruption of the timing or magnitude of these processes can impair scar resolution and compromise regenerative efficacy (7,8,28,29).

Although ERBB4-mediated signaling has been extensively studied in cardiomyocytes, where it promotes proliferation, the contribution to non-myocyte cell populations that shape the regenerative microenvironment remains unknown. The present study therefore investigated the role of ERBB4 in fibroblasts and myeloid cells during neonatal cardiac regeneration.

The central finding of this study is that neonatal deletion of *Erbb4* in fibroblasts or myeloid cells alters the timing and coordination of myocardial repair after MI, while leaving cardiomyocyte mitotic activity unchanged. Fibroblast-specific loss of *Erbb4* resulted in a modest but persistent delay in scar resolution accompanied by transient impairment of microvascular restoration and elevated *Col1a1* gene expression at 10 dpi. In contrast, myeloid-specific deletion elicited an exaggerated early injury response followed by sustained inflammatory and extracellular matrix remodeling activity. Despite a larger early scar burden, scar resolution progressed to levels comparable to controls by 21 dpi, suggesting an adequate compensatory scar resolving response. The overall scarless reparative outcome at 21 dpi remained largely preserved. This observation indicates that neonatal regeneration is remarkably resilient and may rely on abundant compensatory pathways.

### Preserved cardiomyocyte dynamics uncouple the observed phenotypes from cardiomyocyte proliferation

NRG1-ERBB signaling has classically been linked to cardiomyocyte proliferation, with cardiomyocyte ERBB2/ERBB4 activation considered a central driver of neonatal cardiac regeneration (9,11,14,15). Consistent with previous studies, myocardial infarction induced a marked increase in cardiomyocyte mitotic activity. However, pH3⁺ and Aurora B⁺ cardiomyocyte numbers, as well as cardiomyocyte cross-sectional area, remained comparable between WT and both fibroblast– and myeloid-specific *Erbb4*-KO mice. The preservation of cardiomyocyte proliferative and hypertrophic responses represents an important strength of the present models, as it uncouples the observed differences in scar remodeling and repair from alterations in cardiomyocyte proliferation. Consequently, the distinct healing responses can be attributed primarily to ERBB4-dependent functions within fibroblasts and myeloid cells rather than secondary effects on cardiomyocyte proliferation.

### ERBB4 signaling coordinates inflammatory transitions during neonatal cardiac repair

Unlike the adult heart, where MI triggers prolonged inflammation and permanent fibrotic remodeling, neonatal injury elicits a tightly regulated, transient immune response that supports regeneration. Neonatal macrophages promote angiogenesis, debris clearance, and reparative signaling after MI, and their depletion results in persistent scarring. The precision of their temporal activation therefore determines whether injury resolves through by regeneration or through fibrotic remodeling (22,24,30–32). NRG1/ERBB4 signaling, beyond its established role in cardiomyocytes, is an important regulator of immune cell activation and tissue fibrosis. ERBB4 activation has previously been identified as an anti-inflammatory, antifibrotic pathway in macrophages across cardiac, pulmonary, and dermal injury models in adult mice, reducing pro-inflammatory cytokine production and promoting reparative responses (19,21). Our findings extend this to the neonatal regenerative context but reveal a more nuanced role than simple inflammatory suppression.

Myeloid-specific *Erbb4* deletion increased infarct size during the early phase after neonatal MI and resulted in persistent inflammatory pathway activation at 10 dpi together with upregulated *Mmp2/Mmp9* expression. The increased early infarct burden suggests that myeloid ERBB4 contributes to efficient injury containment and tissue remodeling during the acute phase after neonatal MI. At 10 dpi, transcriptomic profiling demonstrated persistent activation of inflammatory, extracellular matrix remodeling, and TGFβ-associated programs. Although such pathways are often associated with ongoing injury responses, their persistence in M-*Erbb4*-KO hearts may also reflect active reparative remodeling. Consistent with this interpretation, increased *Mmp2* and *Mmp9* expression suggests continued extracellular matrix turnover, a process that may facilitate scar resolution and contribute to the normalization of scar size observed by 21 dpi.

Together, these findings indicate that myeloid ERBB4 coordinates the transition from inflammatory activation to reparative resolution rather than simply suppressing inflammation. Despite the altered inflammatory and remodeling profile, myeloid *Erbb4* deletion did not prevent complete scar resolution by 21 dpi, underscoring the robustness of neonatal repair. The final normalization of scar resolution further suggests that regenerative competency is not dictated by a single immune signaling pathway but rather emerges from a network of compensatory mechanisms. Because the LysM-Cre driver targets monocytes and neutrophils in addition to macrophages, the relative contribution of these populations cannot be distinguished in this model. However, ErbB4 expression is induced in activated macrophages but is undetectable or minimal in neutrophils, suggesting that the observed phenotype is predominantly driven by macrophage-specific ERBB4 signaling (21,33).

### ERBB4 in fibroblasts regulates matrix remodeling rather than fibrosis initiation

Fibroblast-specific *Erbb4* deletion revealed a distinct role for the NRG1/ERBB4 pathway in the later remodeling phase of neonatal cardiac repair. Neonatal fibroblasts contribute both to early wound stabilization through transient extracellular matrix deposition and to subsequent scar resolution with restoration of tissue architecture (28,34). Accordingly, fibroblast-specific *Erbb4* deletion selectively disrupted the transition into the reparative remodeling phase, resulting in a limited but significantly larger residual scar at 21 dpi. Collagen deposition in the regenerating neonatal heart follows a biphasic pattern, with an early inflammatory-phase burst followed by a second peak characterized by extracellular matrix remodeling between days 10 and 21 (3,30,35). Persistent *Col1a1* upregulation in FB-*Erbb4*-KO hearts at 10 dpi likely reflects dysregulated entry into this second remodeling phase rather than augmented early fibrosis, implicating ERBB4 in coordinating the fibroblast transition from matrix deposition to resolution. Consistent with this interpretation, early angiogenesis was preserved in both knockout models, whereas FB-Erbb4-KO hearts exhibited only a transient reduction in capillary density at 10 dpi, suggesting that fibroblast ERBB4 contributes to vessel maturation or maintenance rather than angiogenic initiation (36,37).

## Conclusions

We conclude that ERBB4 signaling in fibroblasts and myeloid cells functions as a coordinator of the regenerative microenvironment dynamics rather than as an essential switch for regeneration. By demonstrating that fibroblast-and myeloid-specific *Erbb4* deletion alters the timing and efficiency of inflammatory resolution and extracellular matrix remodeling without affecting cardiomyocyte cell-cycle activity, this study extends the current framework of ERBB4-mediated regeneration beyond cardiomyocytes and identifies non-cardiomyocyte ERBB4 signaling as an important determinant of regenerative repair (6,17,38,39). It supports temporally tuned, combinatorial therapies that restore orderly inflammation and remodeling, rather than simply suppressing them, to limit adverse remodeling and improve functional recovery after myocardial injury. Altogether, our data support the hypothesis that ERBB4 signaling in non-cardiomyocyte cell populations contributes to neonatal cardiac repair through mechanisms distinct from proliferative pathways.

## Author contribution

V.F.M.S, G.W.D.K., and S.V.D.B. conceived and designed research; S.V.D.B., S.E., C.C., E.V.D.B and J.M.T.C. performed experiments; S.V.D.B., S.E. B.K.G. and C.C. analyzed data; S.V.D.B, S.E., G.W.D.K. and V.F.M.S. interpreted results of experiments; S.V.D.B prepared figures; S.V.D.B prepared manuscript. All authors edited, revised, and approved the final version of the manuscript.

## Abbreviations

cTnT: cardiac troponin T
Dpi: days post-injury
ERBB4: Erb-B2 Receptor Tyrosine Kinase 4
FB: fibroblast
FFPE: formalin-fixed paraffin-embedded
KO: knock-out
LAD: left anterior descending artery
MI: myocardial infarction
NRG1: Neuregulin-1
P1: postnatal day 1
P7: postnatal day 7
PFA: paraformaldehyde
pH3: Phospho-histone 3
WT: wild-type

## Acknowledgments

We thank Peter Angel from DKFZ Heidelberg for the kind donation of Tg(Col1a2-cre)23Angl mice. We thank Piet Finckenberg and Päivi Leinikka from the University of Helsinki for surgical training in the neonatal LAD ligation model. We are especially grateful to Eline Roeyen for her expertise and assistance with the western blot experiments. We further thank Tine Bruyns, Mandy Vermont, and Evy Mayeur for their excellent technical support.

